# Categorical and semantic perception of the meaning of call-types in zebra finches

**DOI:** 10.1101/2024.11.14.623689

**Authors:** Julie E. Elie, Aude de Witasse-Thézy, Logan Thomas, Ben Malit, Frédéric E. Theunissen

## Abstract

Vocal communication in social animals involves the production and perception of various calls that ethologists categorize into call-types, based on their acoustical structure and the behavioral context of production. Whether animals perceive these categories and associate distinct meanings to them remains unknown. The zebra finch, a gregarious songbird, uses approximately 11 call-types to communicate hunger, danger, social conflict, and establish social contact and bonding. Using auditory discrimination tasks, we show that the birds discriminate and categorize all the call-types in their vocal repertoire. In addition, systematic errors were more frequent between call-types used in similar behavioral contexts than could be expected from their acoustic similarity. Thus, zebra finches organize their calls into categories and create a mental representation of the meaning of these sounds.

## Main Text

Social animals often possess a rich repertoire of distinct vocalizations used to communicate various needs, internal states or external events, and to elicit appropriate responses in individuals that hear the message (*1, 2*). To understand these vocal signals in terms of a basic code for communication, researchers have categorized the sounds into call-types based on their acoustic properties, the context during which they are emitted, and the responses they elicit in receivers. However, the extent to which animals agree with these human expert’s categorizations remains an open question. Peter Marler, while preforming the first detailed description of the vocal repertoire of the European chaffinch, argued that categorizing vocalizations into types is the first step for understanding the vocabulary that is available for the “language” of a particular species (*3*). Because such classification is performed by human experts based on a multifaceted approach involving behavioral assessments and acoustical analyses, we will refer to the call-types that have been used to organize the vocal repertoire of many species (*3-14*) as ethogram-based call-types or Etho call-types. Our first questions are to establish whether animals can distinguish the calls from distinct Etho call-types, and whether the perception of those communication calls is spontaneously categorical. Previous studies have shown that animals learn to categorize sounds along both natural or arbitrary acoustic dimensions (*15*). They also generalize to novel exemplars in these groupings as long as the categories contain recognizable patterns (*16*). However, whether they have an inherent categorical perception of their own communication calls through normal ontogeny remains an open question.

A second and more profound question is whether animals “understand” the meaning of Etho call-types in a way that matches human interpretations. This is a very difficult question to answer as it would require the researcher to read the animal’s mind. The nature of the animal mind has been an ageless debate, as reflected by the dichotomy between mentalism vs behaviorism schools (*17*). In particular, the differences in the faculty of language in humans versus animals has often been showcased as illustrating the gap in mental representations and cognitive abilities between humans and other animals (*18*). It has remained a challenge to clearly demonstrate that animals’ calls are not simply produced and responded to in a simple reflexive manner. The first demonstration of the potential referential property of animal calls came with the now classic studies of the vervet monkeys’ alarm calls (*19*). Indeed, the monkeys adopt appropriate complex avoidance or escape behaviors in response to the playbacks of distinct alarm calls, without the actual presence of the predator. This functional referential quality has now been demonstrated in multiple mammalian and avian species (*20-25*). However, it remains controversial whether animals hearing a specific alarm call have a mental representation of the predator or reflexively perform the appropriate escape or avoidance behavior that would be triggered upon hearing the specific alarm call. As such, these calls have been labelled as functionally referential, which some argue is fundamentally similar to the general concept of an ethogram-based call-type (*26*). To address the nature of the mental or neural representation of the meaning of call-types, ethologists must therefore rely on indirect measures where observed responses could not simply be explained as actions triggered by specific sound classes but instead by their meaning.

To find-out if animals have a categorical perception of Etho call-types and whether they have a mental representation of the meaning of Etho call-types, we quantified the ability of zebra finches to discriminate all the Etho call-types in their repertoire in a hearing task. The zebra finch is a gregarious songbird that has a rich vocal repertoire used in distinct and well-characterized social behaviors (*7, 8*). Zebra finches can quickly be taught to perform complex acoustical discrimination tasks in operant conditioning paradigms (*27, 28*). Here, we performed such experiments to quantify the birds’ discrimination of all their Etho call-types. Using the birds’ behavioral responses, we first assessed whether this discrimination shows properties of categorical perception, and second whether discrimination performance is sensitive to the putative meaning of Etho call-types beyond what could be expected from acoustics.

## Results

### Zebra Finch Etho Call-Types

The vocal repertoire of the zebra finch has been divided into approximately 11 Etho call-types (*7, 8*), which we further cluster here into six semantic hyper-categories (Fig. 1A). Three Etho call-types form the *contact calls* hyper-category. These calls are used for maintaining spatial cohesion among familiar individuals including pair-bonded partners. The *pair-bonded calls* hyper-category groups two Etho call-types that are produced during the pair-bonding and nest building behaviors. Song is a courtship signal produced by males only and forms its own semantic hyper-category. Zebra finches can also inform each other of danger by emitting two distinct *alarm calls*, which form a fourth semantic hyper-category. The fifth semantic hyper category, the *agonistic calls*, are produced under distress or during fights with conspecifics. Finally, chicks and juvenile birds also produce food *begging calls*, which forms its own semantic hyper-category. One can visualize how distinct the Etho call-types are by projecting the calls of many individuals in a 2D space that characterizes the calls acoustic properties. Such an acoustic space (Fig. 1A, Fig. S1) reveals that the vocal repertoire of the zebra finch is a mixture of discrete signaling, with some Etho call-types found in distinct acoustic clusters (e.g. DC, Be, Te), and graded signaling, with other Etho call-types forming a graded continuum (e.g. Wh, Ne). Despite the differences in the level of clustering, we have previously shown using various types of supervised classifiers operating on distinct acoustic feature spaces that these 11 Etho call-types are acoustically distinct and are correctly classified by algorithms with ∼60% accuracy (Fig. 1B) (*7*).

**Fig. 1.**
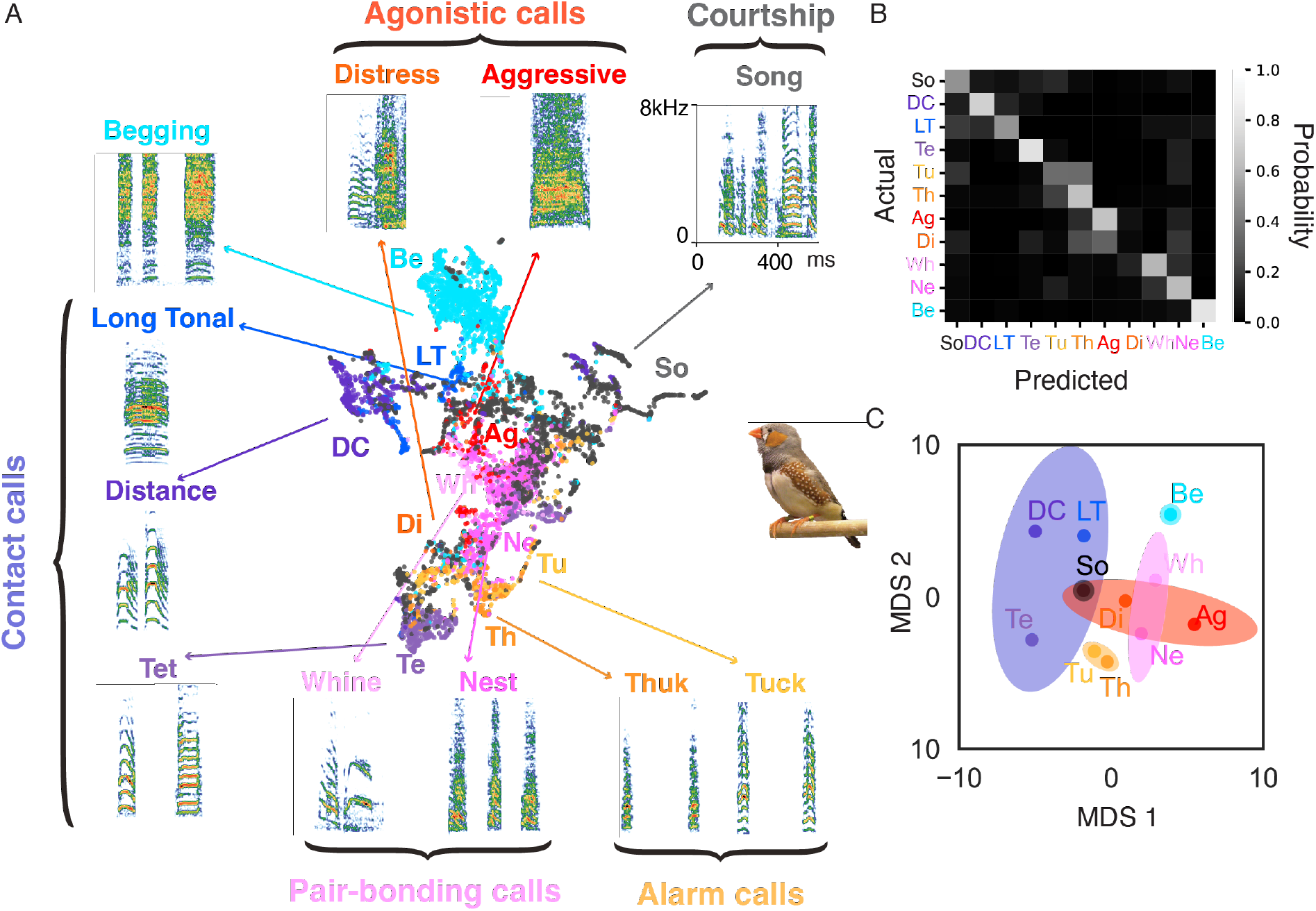
The acoustic space of the zebra finch vocal repertoire. (**A**) The zebra finch vocal repertoire is divided into 11 distinct Etho call-types. ∼8000 call renditions from 45 birds are projected in the two first dimensions of a non-linear embedding (UMAP) applied to predefined acoustic features and color-coded according to the Etho call-type categorization (see methods and Fig. S1). (**B**) Confusion matrix quantifying the cross-validated performance of a supervised classifier (Linear Discriminant Analysis) at classifying calls into the Etho call-types using the spectrogram of each vocalization. The percent correct classification is obtained from the average of the diagonal. Data from(*7*). (**C**) Acoustic distances are derived from the probability of correct and mis-classifications in B (see methods) and used in a multi-dimensional scaling (MDS) analysis to obtain a 2D projection of the acoustic space of ethogram-based call-types. The colored ellipses group call-types belonging to the same semantic hyper-category (Contact Calls: DC, LT, Te; Pair-bonding Calls: Wh, Ne; Alarm Calls: Tu, Th; Non-affiliative calls: Di, Ag; Begging calls; Courtship: Song). The spatial organization of call-types in this acoustic space (based on misclassifications of the spectrogram trained classifier) is contrasted in Fig. 4 to the spatial organization based on behavioral misclassifications (Fig. 2).

### Etho Call-Type Perceptual Discrimination

The fact that animal vocalizations can be hierarchically organized into Etho call-types and semantic hyper-categories based on joint behavioral observations and acoustic properties does not tell us whether animals can themselves discriminate between Etho call-types and whether they, perceptually, also group Etho call-types using a hierarchical organization. To evaluate how adult zebra finches (6 males and 6 females) discriminate the Etho call-types of their repertoire, we tested their abilities to discriminate one Etho call-type from all others in an auditory Go-NoGo task, systematically going through all 11 Etho call-types (Fig. 2A)(*27, 28*). Each Etho call-type corresponded to a set of vocalizations produced by a dozen different birds (see Methods). Since the same stimulus (same rendition from the same vocalizer) was rarely heard twice (see Methods), this discrimination had to rely on the acoustic features that are invariant and specific to each Etho call-type. Figures 2B and 2C show the learning curves of one female and one male for the first two Etho call-types and the last Etho call-type that they were tested with. Every one to three days, the birds are asked to learn a new reward contingency (a new Etho call-type is the rewarded set of calls). Despite this, zebra finches could relatively quickly learn to categorize this heterogenous set of call stimuli into Etho call-types. Almost all birds succeeded in quickly discriminating all the 11 Etho call-types (Fig. 2D; 127/131 Fisher Exact tests (12 birds*11 call-types – 1) on the probability of correct decision with p<0.05). Only two birds failed to discriminate Ag calls, a single bird failed to discriminate Di calls and another single bird failed to discriminate Ne. Because our experimental design aimed at quantifying rapid classification performance, it is very likely that for the four failed tests, birds could have performed these specific discriminations given more testing time. Despite these very few failures, the probability of correct classification averaged across all birds and Etho call-types was highly significant with a large effect size (Likelihood ratio test (LRT) on *reward contingency* in the Generalized Linear Mixed Effect (GLME) model comparison; χ^2^(1) = 31.125, p<2.4 10^−18^, Odds Ratio for Reward Contingency, log_2_(*OR*_*RC*_) = 2.47 [2.06, 2.88] 95%; see methods). Male and female birds performed equally well (LRT on *bird subject sex* in the GLME model comparison, χ^2^(2) = 1.79, p=0.41). Although performance varied across Etho call-types (LRT on *tested call-type* in the GLME model comparison, χ^2^(20) = 356.11, p<2.2 10^−16^), all Etho call-types were discriminated above chance level (see Table S1). Thus, Zebra finches excel at discriminating all the Etho call-types of their vocal repertoire.

**Fig. 2.**
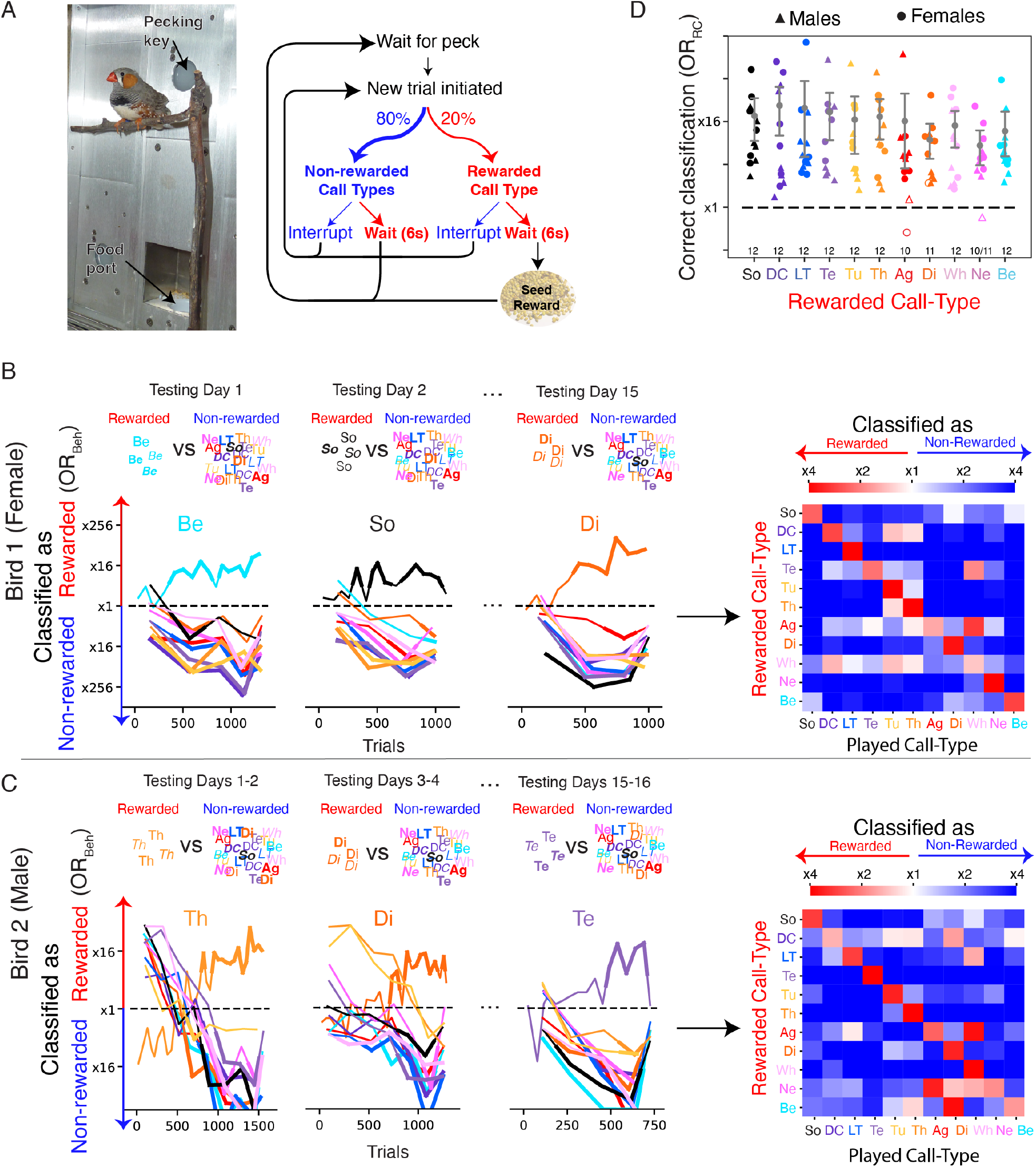
Behavioral discrimination of Etho call-types. (**A**) Zebra finches are trained in an operant conditioning task to interrupt non-rewarded stimuli and to refrain from interrupting rewarded stimuli (*27, 28*). Rewarded stimuli are random samples of one particular Etho call-type and non-rewarded stimuli are random samples taken from all other Etho call-types. Stimuli are sampled across many renditions (>8000) from multiple male and female bird vocalizers (n=45). (**B**) and (**C**). Performances of two birds (B, female; C, male) in this Etho call-type discrimination task. For a given call-type test, birds have up to three consecutive days to identify the rewarded Etho call-type and show their discrimination performance. The first two tests and the last test are shown here. All tests for these two birds can be seen in Fig. S2 and S3. The learning curves show for each Etho call-type (line color) the average bird performance at classifying calls as rewarded or non-rewarded stimuli. The performance is quantified by the odds ratio of correct behavioral classification, *OR*_*Beh*_ (see Methods). Chance performance is *OR*_*Beh*_ ≈ ξ1. Bold lines indicate a significant classification performance (Fisher Exact Test, p < 0.05). The confusion matrices show the classification performance obtained on the last day of each call-type test. (**D**) The performance at classifying the rewarded Etho call-type on the last day of each call-type test is quantified by the odds ratio of stimulus interruption following the reward contingency, *OR*_*RC*_ (see Methods). These values are similar to the diagonal of each subject confusion matrix shown in B and C. Birds for which *OR*_*RC*_ is significantly greater than x1 are indicated by filled symbols while open symbols indicate non-significant tests. The total number of significant tests is indicated at the bottom of the plot (Fisher Exact Test, p < 0.05, n=12 except for Ne where n=11). The grey circles and error bars are the mean and the 95% confidence intervals obtained by bootstrap across birds.

### Etho call-type Categorization

To perform the vocalization discrimination task described above, birds needed to form categories based on invariant features specific to each Etho call-type. In that experimental design however, it remained unclear if birds were only learning the contingency rule (which Etho call-type was rewarded for each test), or if they were also learning the invariant acoustic features that were specific to each category without referring to any internal categorical perception of these Etho call-type. Indeed, because the Etho call-types were also systematically assigned to a specific reward contingency, one could argue that birds were incited to quickly learn the acoustic features of rewarded and non-rewarded categories, albeit using an appropriate generalization rule for novel exemplars (*16*). To more directly test the perception of Etho call-types as categories, we designed a second experiment where birds would only perform well if they were to learn the exact acoustic features of the stimuli in each category and would partly fail if they were to follow the Etho call-type categorization rule.

In this new series of discrimination tests, zebra finches had to classify two groups of vocalizations that were successively congruent and incongruent with Etho call-type categories (Fig. 3). On a first learning day, zebra finches (n=7) were trained to discriminate two Etho call-types, either DC vs Te or Th vs Te, that had been recorded from four different vocalizers. All birds easily performed that simple discrimination task (Fig. 3A and 3B left panels; 7/7 Fisher Exact Test with p<0.05; LRT on *reward contingency* in the GLME model comparison: χ^2^(1) = 21.97, p=2.77 10^−6^, log_2_(*OR*_*RC*_) = 5.61 [4.82, 6.40] 95%). On the following day, the birds were asked to discriminate the Etho call-types of these same vocalizers (Old) and of four new vocalizers (New) using the same reward contingencies (congruent test on Day 2; Fig. 3A and 3B middle panels). As shown in Figure 3D, and in accordance with our results on Etho call-type discrimination, all birds performed well above chance level on the discrimination of calls from the new vocalizers (7/7 Fisher exact test with p<0.05, LRT on *reward contingency* in the GLME model comparison: χ^2^(1) = 18.41, p=1.78 10^−5^, log_2_(*OR*_*RC*_) = 4.86 [3.81, 5.92] 95%). The Etho call-type discrimination performances were not different between calls from new and old vocalizers (Fig 3D ; Old vocalizers calls: 7/7 Fisher exact test with p<0.05, LRT on *reward contingency* in the GLME model comparison: χ^2^(1) = 20.7, *p* = 5. 36 10^−(^, log_2_(*OR*_*RC*_) = 5.4 3 [4.40, 6.28] 95%;

**Fig. 3.**
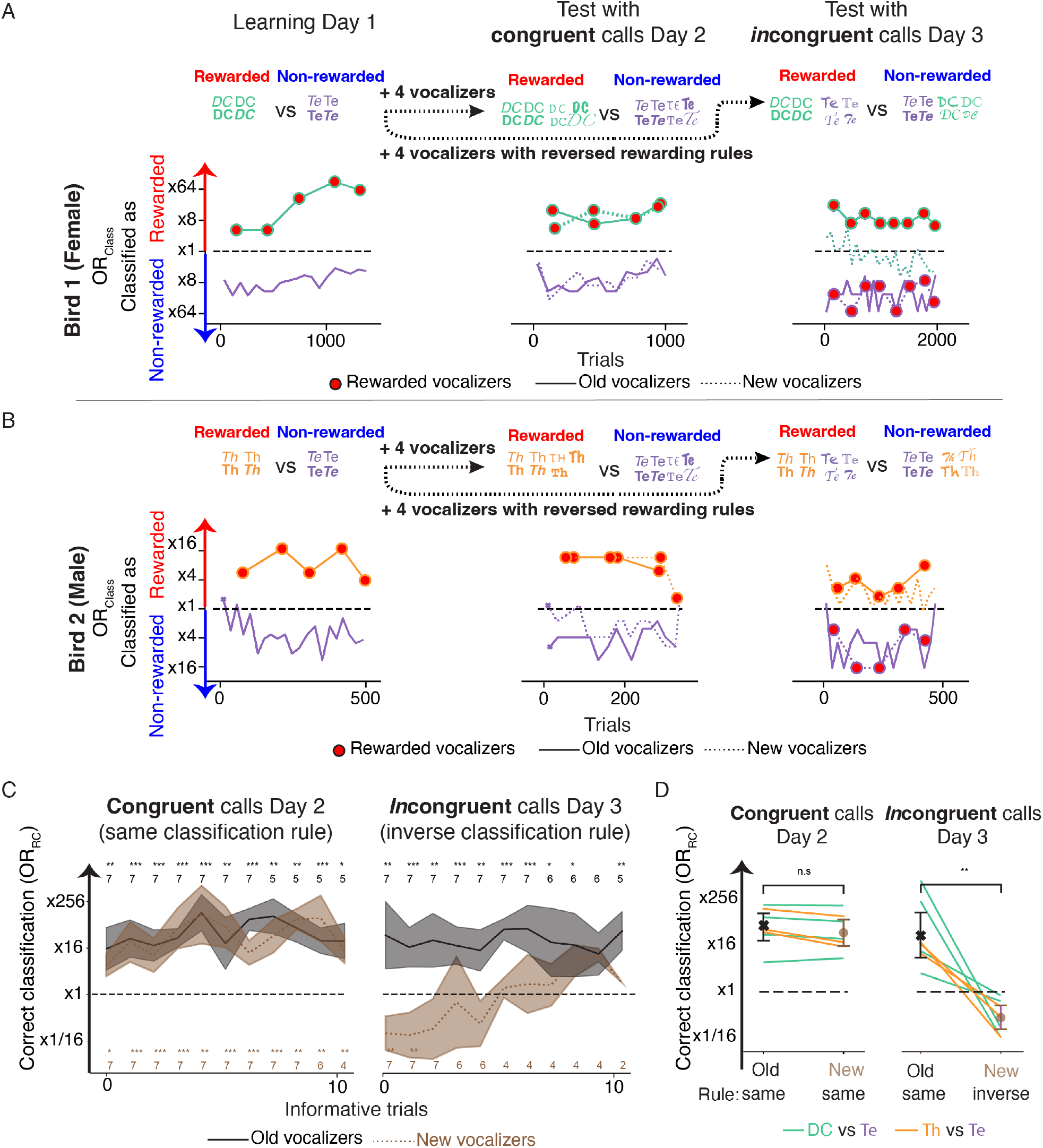
Categorical perception of Etho call-types. (**A**) and (**B**) Classification performances of two example subjects (A, female; B, male) in the categorical perception experiment. Zebra finches are trained to discriminate between two call-types on Day 1. Their ability to generalize to new vocalizers is tested by adding calls from four new vocalizers on both Day 2 and 3, and inversing the rewarding rule with respect to Etho call-type for the vocalizers added on Day 3 (incongruent calls) but not on Day 2 (congruent calls). The learning curves show for each Etho call-type (color) and each set of four vocalizers (Old, solid line; New, dashed line), the average bird performance (Odds Ratio for Classification, *OR*_*Class*_, see Methods) at classifying calls as rewarded or non-rewarded stimuli. Chance performance is *OR*_*Class*_ ≈ ξ1. (**C**) Evolution of the average performance of all birds (n=7) with the number of informative trials (i.e. the trials that are uninterrupted and thus provide feedback on the reward contingency, see Methods). Since the number of informative trials was variable across subjects, the exact number of subjects for each data point is displayed at the top and bottom of the graph. The performance is quantified by the odds of correctly classifying calls according to reward contingency *OR*_*RC*_. Statistical significance is calculated using a LRT on *reward contingency* in a GLME model comparison (see Methods). (**D**) Average performance of all birds (n=7) for Old versus New vocalizers on the congruent and incongruent test days. * p<0.05, **p<0.01, ***p<0.001.

LRT on *New vs Old* in the GLME model comparison, χ^2^(2) = 3. 34, *p* = 0.188). Impressively, birds immediately categorized the calls from the new vocalizers correctly (Fig. 3A and 3B middle panels and Fig. 3C, congruent). To quantify this effect, we examined the early behavioral responses on that day, before the birds could acquire information on reward contingency for all vocalizers and call-types. On these trials, where birds could only rely and apply the reward contingency rule learned on the previous day, the birds correctly classified the Etho call-types of the new vocalizers they had never heard with high accuracy (New vocalizers data, LRT on *reward contingency* in the GLME model comparison: χ^2^(1) = 6.12, *p* = 0.01 34, log_2_(*OR*_*RC*_) = 3.22 [1.06, 5.37] 95%). Their performance was also not significantly different from that of old vocalizers’ calls (Old vocalizers data, LRT on *reward contingency* in the GLME model comparison: χ^2^(1) = 7.21, *p* = 0.0072, log^2^(*OR*_*RC*_) = 4. 3 3 [1.26, 7.40] 95%; all vocalizers data, LRT on *New vs Old* in the GLME model comparison: χ^2^(2) = 0.71, *p* = 0.701). These values of classification performance remained similar throughout the day (Fig. 3C, congruent): birds easily generalized the classification rule of Etho call-types to new vocalizers.

On the third day of the experiment, the birds were again asked to discriminate the same Etho call-types from yet another set of four new vocalizers, but this time, the calls from these new vocalizers were following the opposite reward contingency to the calls from the old vocalizers (incongruent condition; Fig 3A and 3B right panels). Consistent with the hypothesis that the birds were performing a call-type categorization, we found that both the old and the new vocalizers’ calls were classified using the reward contingency from the previous days: calls from new vocalizers lead to systematically wrong decisions for getting rewards resulting in a flip of the value of the log_2_(*OR*_*RC*_) (Fig 3D, incongruent). All birds classified the congruent calls according to the reward contingency rule well above chance level (7/7 Fisher exact test with p<0.05; LRT on *reward contingency* in the GLME model comparison: χ^2^(1) = 10.7 35, p = 0.00105, log^2^(*OR*_*RC*_) = 4.84 [2.92, 6.76] 95%). In contrast, the incongruent calls were significantly *below* chance level, showing that the birds failed to follow the inverse reward contingency rule of these new calls (4/7 Fisher exact test with p<0.05; LRT on *reward contingency* in the GLME model comparison: χ^2^(1) = 8.81, *p* = 0.0029 3, log_2_(*OR*_*RC*_) = –2.17 [– 3.21, –1.12] 95%). As a result, the classification performance according to the reward contingency was different between calls of new and old vocalizers (LRT on *Old vs New* in GLME model comparison: χ^2^(2) = 1654, *p* < 2.2 10^−16^). Thus, birds continued to apply the discrimination rule that they learned on prior days, classifying the new calls based on their Etho call-types and not based on their reward contingency. This effect was particularly clear from the responses gathered up to the first informative trials, before the birds could acquire information on reward contingency for all vocalizers and call-types (Fig 3C, incongruent; New: log_2_(*OR*_*RC*_) = – 3.57 [–5.25, –1.88] 95%; Old: log_2_(*OR*_*RC*_) = 5. 36 [1.91, 8.80]95%; LRT on *Old vs New* in GLME model comparison: χ^2^(2) = 104.6 3, *p* = 1.9 10^−23^). Over the day, the birds slowly learned to flip the reward contingency for the new vocalizers’ calls, indicating that they could learn to recognize some discriminable acoustic features in these calls and apply the new arbitrary reward contingency rule. We have previously shown that zebra finches excel at recognizing the individual signature of vocalizers for any call-type (*27, 28*). Thus, the failure of correct classification in the incongruent condition is unlikely due to the absence of discriminable acoustic information, but rather, demonstrates that zebra finches trained at Etho call-type discrimination, will spontaneously group novel vocalizations using Etho call-types and that, in this context, this categorization hinders the learning of other potentially behaviorally relevant categories, such as the ensemble of calls produced by the same individual.

### Hierarchical Perception of Call-types Based on their Meaning

Next, we quantified the extent to which the acoustic features and acoustic clustering properties of vocalizations could explain the birds’ behavioral categorization of calls into Etho call-types. We also asked if zebra finches could confuse call-types that were semantically related but acoustically distinct, providing evidence of an internal model of Etho call-types based on their meaning.

The first step was to establish a measure of acoustic distance between Etho call-types that we could compare to a measure of behavioral perceptual distance. The acoustic discriminability of the Etho call-types can be quantified by a supervised classifier operating on acoustic features that describe the sounds (*7*). The classifier (Linear Discriminant Analysis, LDA) is trained to categorize calls along the Etho call-types and is used to predict the Etho call-types of novel calls. These predictions are gathered in a confusion matrix showing the conditional probabilities of Etho call-type prediction given the actual Etho call-type (Fig. 1B). The diagonal of this matrix shows the average probability of correct classification for each Etho call-type (Mean = 56± 4 %, 2SE bootstrap) and the off-diagonal terms quantify systematic errors of the classifier. For example, the classifier often confuses calls belonging to the same hyper categories, such as the alarm calls (Tu and Th), the agonistic calls (Di and Ag), and the bonding calls (Wh and Ne). These miss-classification probabilities can be interpreted as acoustic distances between pairs of Etho call-types (Acoustic confusion matrix, Fig. 4A; see Methods) and can be compared to the measures of perceptual distance obtained from the behavioral discrimination performance of birds in the first series of behavioral experiments (data from two example subjects in Fig. 2B and 2C; Perceptual confusion matrix, Fig. 4A; see Methods).

**Fig. 4.**
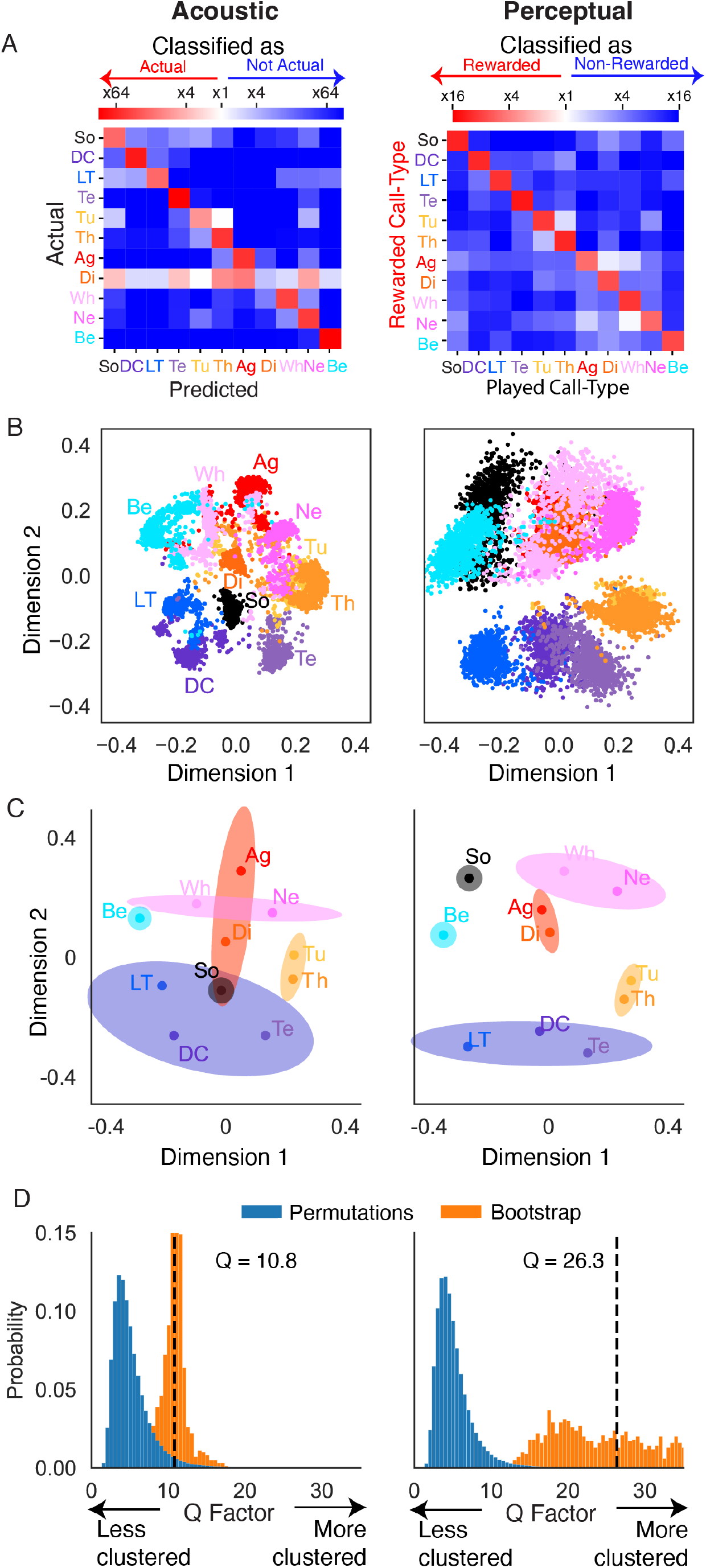
Acoustic and perceptual space of Etho call-types. (**A**) The algorithmic and behavioral performance of classification of all Etho call-types are compared to evaluate if birds’ misclassifications are better explained by semantic rather than by acoustic similarities. In the acoustic space, the confusion matrix of classifier performance (*OR*_.*C*_, left) is obtained from the matrix of classification probabilities (Fig. 1B, see methods). In the perceptual space, the confusion matrix of behavioral performance (*OR*_*Beh*_, right) is the average performance of all subject birds in the Etho call-type discrimination experiment (Fig. 2; 6 males, 6 females). (**B**) 2D projections of Etho call-types generated by applying multi-dimensional scaling (MDS) to the distances given by the off-diagonal values (systematic misclassifications) of each of the two matrices in A. The clouds of points are obtained by bootstrapping across vocalizers in the call database (Acoustic, n=45) or subjects in the behavioral experiment (Perceptual, n=12). (**C**) Centroids (mean) of each cloud in B are shown as well as illustrative ellipses grouping the Etho call-types into the semantic hyper-categories described in Fig. 1. Note that to facilitate the visual comparison of the 2D acoustic space and 2D perceptual space of Etho call-types, a Procrustes transformation (scaling and rotation) was applied to the acoustic space (see Fig. 1C for the original projection). (**D**) Histogram of Q factor (ratio of the between to within hyper-category Euclidian distance, see Methods) used to quantify the clustering of Etho call-types into semantic hyper-categories. To assess significance, a null distribution of Q values based on sampling random hyper-categories of same size was generated by permutation (Perm, blue bars). To assess confidence levels on the actual Q value measured for the actual semantic hyper-category a bootstrap was performed on the acoustic dataset across vocalizers (n=45) and on the behavioral data set across subjects (n=12, Boot, orange bars). X-axis is truncated at Q=35.

To visualize and compare these acoustic and perceptual distances between pairs of Etho call-types, we applied a multi-dimensional scaling analysis (MDS) which produces 2D maps displaying the position of Etho call-types in acoustic or perceptual spaces (Fig. 1C, 4B and 4C). The variability in the calculations and robustness of these maps are evaluated by bootstrap procedures across birds (Methods) and are displayed in Figure 4B. It is apparent from visual inspection that the acoustic and behavioral maps are similar. This match is quantified by a low value of Procrustes disparity, a measure of the mean error of the match (Proc. Disp. = 0.042 ± 0.023 2SE bootstrap; normalized effect size: R^2^ = 0.703, p=0.0014, permutation test). Thus, behavioral performance in Etho call-type categorization and their systematic misclassifications can in part be attributed to the acoustic distances between calls.

However, one can also observe in Figure 4C that the Etho call-types belonging to the same semantic hyper-category, i.e. that are closer in meaning, are closer in the perceptual space than they are in the acoustic space. This merging of Etho call-types in the perceptual space indicates that birds perceptually misclassified these calls more often than expected from their acoustic distance. To quantify the clustering effect of Etho call-types into semantic hyper-categories, we measured, in each map, how close each Etho call-type is on average to the semantic hyper-category it belongs to, as compared to other semantic hyper-categories (distance ratio Q). By the definition of Q, the Etho call-types are on average Q times closer from the centroid of the semantic hyper-category they belong to, than to the centroids of other semantic hyper-categories. In other words, a higher value of Q indicates a stronger clustering of Etho call-types along the semantic hyper-categories. In both the acoustic and perceptual maps, values of Q were higher than expected for random hyper-semantic groupings obtained in a permutation test, validating the semantic hyper-categories both in the acoustic and perceptual spaces (perceptual map: Q = 26.35 ± 17.49 2SE; p=0.0019; acoustic map: Q = 10.82 ± 3.14 2SE; p=0.0376). Consistent with the merging of Etho call-types in the perceptual space visible in Figure 4C, the grouping of semantically related Etho call-types was ∼2.43 times greater in the perceptual space than in the acoustic space (Q is larger in the perceptual map than in the acoustic map, p=0.004, bootstrap). Thus, zebra finches make systematic errors in call-type categorizations with similar meanings at much higher rates than what would be expected from their acoustic distances. Borrowing from a term coined by Patricia Kuhl to describe the categorical perception of phonemes (*29*), we call this local shrinkage of the perceptual space centered on semantic hyper-categories of Etho call-types, the *semantic magnet effect*.

## Conclusions

We have shown using operant conditioning that zebra finches are able to discriminate all the Etho call-types in their vocal repertoire. They learn to perform this categorization task rapidly and generalize to novel stimuli from renditions of vocalizers they have not heard before. Moreover, these natural category boundaries appear to hinder the learning of other categories such as unique vocal signatures. Finally, the analysis of the systematic errors performed during attempted discrimination shows that zebra finches mistake call-types from similar semantic hyper-categories more often than expected from their acoustic similarities: the semantic magnet effect.

In multiple animal species, ethologists have described Etho call-types without gathering evidence that animals would agree with their repertoire organization. In the case of alarm calls, such as those used by vervet monkeys (*19-21*) or other animals including birds (*30, 31*), the sophisticated usage and appropriate complex behavioral responses elicited in the receivers clearly demonstrated the specificity of both the information sent and the adapted behavioral responses. The fact that these alarm call-types can be interpreted within the broad context of alarm responses to elicit behaviors that are specific to each type of predator and to the level of danger show that indeed these Etho call-types are discriminated by animals. Nonetheless, until now, there had not been a single study that attempted to assess whether all the Etho call-types comprising a vocal repertoire of a particular non-human species could be acoustically discriminated by the members of that species. Drawing on previous characterization of the zebra finch vocal repertoire (*7, 8*), we leveraged a behavioral task design where we could test all combinations of Etho call-types discrimination and demonstrated that this discrimination is indeed performed by zebra finches for the 11 Etho call-types of their repertoire. Furthermore, this trained discrimination shows generalization and robustness that is indicative of a form of categorial perception. Just as humans discriminate isolated vowels (*32*) and marmosets categorize some of their call-types (*33*), zebra finches do not only categorize call-types that are clearly distinct acoustically, but also those found along a graded acoustical space such as the one occupied by whine and nest calls, or tet and thuk calls (Fig. 1). Thus, categorical processing of meaningful categories, species-specific call-types, is not specific to primates.

Showing that animals understand the vocal messages in an anthropomorphic sense is difficult because of the relative invariability of natural contexts of production and of the specificity of appropriate behavioral responses. A reflexive mechanism with a one to one mapping between the acoustic characteristics of the sound perceived and the response could be the simplest explanation. A “deeper” understanding from the part of the receiver would imply the presence of a mental representation of the message that could be used flexibly. And indeed, here, we showed that zebra finches can flexibly respond to calls in their repertoire and succeed in an auditory discrimination task of Etho call-types without reflexively responding to these calls. Hearing a call does not necessarily require a stereotypical behavioral response, such as escaping when hearing alarm calls.

Direct evidence for the birds’ mental states when hearing their conspecific calls is impossible, but indirect evidence can be gathered. For example, it has been shown that when Japanese tits are primed with species specific alarm calls for snakes, they scan their visual space and perform investigative behaviors to wood sticks that are moved in snake like motions (*34*). A simple interpretation of these results is that the snake alarm-call elicited a mental imagery of a snake, which yielded unexpected and surprising responses to moving sticks. In the present experiment of Etho call-type discrimination, zebra finches confuse Etho call-types that have a similar meaning at a higher rate than would be expected based on the acoustic similarity of these call-types. This semantic magnet effect could be explained by postulating that the playback of calls during the task elicits mental representations based on the meaning of calls. From a mechanistic perspective, the representation of Etho call-types in higher-order auditory brain regions would perform a non-linear mapping of the acoustic information. The representation in the higher auditory centers would be driven or organized by the usage or “meaning” of the Etho call-type. For example, neural responses recorded in the pre-frontal cortex of rhesus monkeys have shown to be organized based on semantics over acoustics (*35*). Note that these *semantically warped neural representation* could underlie, or not, mental representations of meaning. In the zebra finch, we have shown that Etho call-types can be correctly decoded from the neural responses of avian neurons in cortical-like auditory pallium (*36, 37*). It remains to be seen to what extent these neural representations participate in the behavioral discrimination of Etho call-types used during natural communication, and to what extent they are shaped by the semantic similarities: can we define distances in the neural space that also display the magnet effect? Whereas a coarse semantic cortical map has been described in humans listening to stories (*38*), such neural correlate has not yet been found in non-human animals.

Finally, two of the trademarks of categorical perception and in particular of the perceptual magnet effect for phonemes in humans are that it is, in part, acquired during development by exposure (*39*) and that early exposure can have a lasting effect (*40, 41*). It is also postulated that phonemic categorical boundaries can be learned prior to and for facilitating language acquisition (*39*). Similarly, it has been shown that the interpretation and usage of appropriate call-types in non-human primates can be in part dependent on exposure and usage (*42*). In songbirds, there is a very rich body of literature that convincingly shows the crucial role of sensory exposure to song for normal song production (*43, 44*) and for song selectivity in higher auditory areas (*45, 46*). Research in the ontogeny of both the production and perception of all Etho call-types in the zebra finch could provide further proof for the semantic magnet effect and its neural underpinnings.

Although the extent of the semantic quality of animal vocal communication signals should be debated (*20, 26*), common themes in the use of complex acoustic signals and their neural mechanisms are abundant particularly in birds. Avian call-types have been shown to be functionally referential (*21*), used in combinations to form rudimentary syntax (*47*), capable of eliciting visual imagery (*34*), under volitional control (*48*), deceitful (*49*), and, as demonstrated here, organized perceptually according to their meaning. Thus, avian communication offers unique opportunities to investigate the neural mechanisms in vertebrates that are essential for producing and interpreting communication calls (*43*). Its study promises to bring invaluable insights into the evolution of vocal communication systems (*50, 51*) and intelligence (*52*).

## Author contributions

Conceptualization: JEE, FET

Formal analysis: FET

Methodology: JEE, LT, FET

Investigation: JEE, AdWT, BM, LT

Software: JEE, AdWT, LT, FET

Resources: FET

Visualization: JEE, AdWT, LT, FET

Funding acquisition: FET

Project administration: JEE, LT, FET

Supervision: JEE, LT, FET

Writing – original draft: FET

Writing – review & editing: JEE, AdWT, BM, LT, FET

## Data and materials availability

All behavioral data and the code used in the analysis is publicly available on Zenodo at https://zenodo.org/records/13623969.

The acoustical database of calls is publicly available in Figshare at https://doi.org/10.6084/m9.figshare.11905533.v1.

## Supplementary Materials

Materials and Methods

Figs. S1 to S3

Tables S1 to S2

References 53

