## Supplementary Materials for "Categorical and semantic perception of the meaning of call-types in zebra finches"

**The PDF file includes:**

Materials and Methods
Figs. S1 to S3
Tables S1 to S2

### Materials and Methods

Ethic Statement.

Animal experiments were approved by the Institutional Animal Care and Use Committee (IACUC) of the University of California, Berkeley (US; UCB protocol number: AUP-9157) following the guidelines of NIH and AAALAC.

Etho call-type database.

We used the zebra finch call database previously recorded and analyzed in our laboratory (*7, 53*). This vocalization database is composed of 8,136 calls and song syllables from 45 birds (27 adults + 18 chicks: 20 females + 23 males + 2 unsexed chicks). All the birds were born in captivity and housed in the Theunissen Lab zebra finch colony at UC Berkeley. In the colony, birds are bred and housed in large aviaries containing 1 to 15 families and can see and vocally interact with the rest of the colony. The database contains calls from all the call-types in the zebra finch vocal repertoire. The calls were classified into call-types by human experts based on the behavioral context during which they were produced and their acoustic quality. For this reason, we denote this call-type classification ethogram-based (Etho call-types). The Etho call-types include the begging call produced by chicks for feeding (Be); three contact calls: the loud distance call (DC) and the soft Tet (Te) produced by adults and the long-tonal call (LT) produced by chicks; two pair-bonding calls produced during nest building and other sexual behaviors: the nest call (Ne) and the whine call (Wh); two non-affiliative calls produced during aggressive encounters: the aggressive call (Ag) and the distress call (Di); two alarm calls: a general alarm called the Tuck (Tu) and an alarm call directed to chicks called the Thuk (Th); and finally, the song (So) produced by adult males only as a sexual display signal. Although this classification covers all the calls in the repertoire, other classifications, based on acoustic quality only, can use a more refined (e.g. including stack, hat and sweeps in Te) or slightly different partitions. See (*7*) for additional details.

Operant Conditioning for Etho Call-Type Discrimination (Task 1) and Categorization (Task 2).

*Task design*

Detailed description of the operant task and apparatus can be found in (*27, 28*). In brief, each bird subject is placed in an operant chamber set up with a speaker, food hopper, water bowl, and backlit pecking key (Med Associates). The task consists in discriminating between two sets of calls, one of which is rewarded by a 10 second access to seeds (Re stimuli), while the playback of the other set of calls is free of consequence and not rewarded (NoRe stimuli). The bird initiates trials by pecking on the backlit key, which triggers a 6 second stimulus playback, after which a reward is given by raising the food hopper for 10 s if the stimulus is a rewarded one (Re trial). Alternatively, the bird may peck the key at any time during the 6 s playback period to terminate the trial and begin a new trial with a random stimulus. In this case, no food reward will be given regardless of the nature (Re or NoRe) of the stimulus which playback was interrupted. By design, 20% of trials are rewarded while 80% of trials are not rewarded (Fig. 2). To maximize the rate at which reward is received in a session, the bird learns to promptly initiate a new trial and effectively interrupt stimuli that are recognized as NoRe to avoid the full waiting period and move on to the next trial. A bird discrimination is measured by the difference in the proportion of Re and NoRe stimuli that are interrupted. Birds maximize their reward intake by interrupting NoRe stimuli and refrain from interrupting Re stimuli. A quantitative analysis of response times showed that zebra finches have a tendency to double peck. This double pecking is clearly evident on the histogram of response times that showed a very clear bimodal distribution with peaks at 100 and 900 ms with a clear trough at 500 ms. For this reason, we ignored the trials with interruption times below 500 ms.

The presentation of the sound stimuli, the detection of key pecks, and the operation of the food hopper were controlled by a Python program. We used a custom branch of the Python-based pyOperant software (https://github.com/theunissenlab/pyoperant), originally developed by J. Kiggins and M. Thielk in T. Gentner's laboratory at University of California San Diego (https://github.com/gentnerlab/pyoperant).

Between daily sessions, subject birds are housed together in group of two to four birds. Their motivation to work for food is maintained by restricting their food access while maintaining a free access to water. Birds weights are tightly monitored twice a day before and after performing in the operant chamber and, if necessary, their daily seed consumption in the operant chamber is complemented at the end of the day to 3g to ensure that their weights is not dropping below 10% of their initial weight. Once trained, birds were able to get their complete daily food allowance (3g/day) during the testing period.

*Shaping*

The birds learn to use the apparatus during a shaping period that lasts approximately one week. After a short period of acclimatation to the chamber and to the feeder, the birds learn to associate the action of pecking the key with the playback of rewarded (Re) stimuli followed by a short access to seeds in the feeder. Then, the playback of non-rewarded (NoRe) stimuli is progressively introduced and their proportion increased to 80% of the number of playbacks. At this stage, birds are expected to learn to interrupt the playback of NoRe stimuli to more quickly initiate a new trial and optimize their access to the feeder. The shaping was slightly different for the two distinct tasks that birds were then assigned to. In the case of the call-type discrimination task (Task 1), the stimuli chosen during this shaping period were two distinct call-types (CT1 and CT2) randomly chosen for each subject from the 11 Etho call-types in the repertoire. A bird was considered trained on using the apparatus once it had successfully performed above chance level three successive discrimination shaping tests: CT1 as Re stimuli *vs* CT2 as NoRe stimuli; CT2 as Re stimuli *vs* CT1 as NoRe stimuli; CT2 as Re stimuli *vs* all other call-types as NoRe stimuli. 14 out of the 16 birds that were trained to use the apparatus for Task 1 passed the shaping criteria of Task 1. In the case of the categorization task (Task 2), the stimuli chosen during the shaping period as Re and NoRe stimuli were two clearly distinct songs then two clearly distinct call-types (DC vs Tet or Thuk vs Tet). A bird was considered trained on using the apparatus once it had successfully performed above chance level the two discrimination shaping tests. Eight out of eight birds passed the shaping criteria of Task 2. For all shaping tests, the daily performance of the bird subject was measured by the difference in interruption rate of NoRe stimuli as compared to Re stimuli. The statistical significance was estimated at the end of each day using an exact Fisher test.

*Testing*

Once the birds learned to use the apparatus, we tested how they classify their Etho call-types in two distinct tasks: a call-type discrimination task (Task 1) and a categorization task (Task 2). These tasks where performed in two separate sets of birds. Although we labeled Task 1 as discrimination and Task 2 as categorization, both tasks require generalization across vocalizers and renditions (see also Stimulus Design for Call-Type Categorization and Discrimination, below). In Task 1 the need for categorization is implicit, whereas it was explicitly tested in Task 2.

For Task 1, we tested 12 adult zebra finches (6 males and 6 females) in a one (Re) vs all-other (NoRe) call-type discrimination. Two additional birds (1 male and 1 female) were also tested in this task but were removed from this dataset because equipment failures required us to increase the number of testing days for particular call-types which could affect the results. The 12 birds were tested on all combinations of one vs all-other call-types on successive days; the single rewarded call-type was changed to systematically span all call-types in the repertoire and the sequence order was randomized across subjects. For each comparison, the subject had up to three days to identify the rewarded call-type and correctly discriminate it against other call-types. Correct discrimination was assessed at the end of each day by a Fisher exact test based on the number of interrupted trials for Re vs NoRe. The reward contingency was changed to the next Re call-type on the following day if the bird showed a significantly higher number of interrupted trials for the NoRe relative to the Re stimuli or if significance was not reached after three days of successive testing with the same Re call-type. The 3-day limit was set so that the test had a reasonable time limit of 33 test days (11 call-types x 3 days). However, many birds only needed a single test day to perform the discrimination for most of the call-types. Across birds, the median number of days to perform Task 1 (11 call-types) was 19 days with a range of 11 to 31 days. Zebra finches trained on this task initiated a high number of trials each day (number of trials per bird: mean, 11.1; range, 6.8 to 21.8). Across all birds, the total number of testing days was 245 with a total of 132,838 trials (~542 trials per day).

For Task 2, we tested eight adult zebra finches (four males and four females) in an explicit call-type categorization task. The task consisted of one day of training and two days of testing. On the first day of learning, the birds learned to discriminate between two call-types (Te vs Th or DC vs Te) based on distinct renditions from four vocalizers, referred to as “old” vocalizers on subsequent days (see Table S2). On the two testing days, we then explicitly assessed how birds generalized the call-type discrimination to four additional new vocalizers, and either respected the reward contingency assigned on Day 1 (test performed on Day 2) or reverse the reward contingency for these new vocalizers (test performed on Day 3). Note that the new vocalizers introduced on Day 2 and Day 3 were distinct birds (eight new vocalizers total) that the subject had never heard before. Because the reward contingency of the new vocalizers on Day 2 is the same as the one learned on Day 1 for the old vocalizers (same call-type is rewarded), we call this condition *congruent*. On the other hand, because the reward contingency of the new vocalizers on Day 3 is the opposite to the one learned on Day 1 for the old vocalizers (what is the non-rewarded call-type for old vocalizers is the rewarded one for new vocalizers), we call this condition *incongruent*. For example, if a bird had learned on Day 1 to discriminate Te vs DC, where Te was the rewarded call-type, then on Day 3, the Te*s* from the previously learned four vocalizers remained rewarded but the Te*s* from the four new vocalizers were *non-rewarded*, and it was the DCs from the four new vocalizers that were rewarded. In the analysis, we asked whether the birds would continue to classify the stimuli from the new vocalizers using the call-type categorization learned on previous days or whether they would base their discrimination on the idiosyncratic features found in the calls of each vocalizer (*27*). Of all possible pair-wise comparisons, we chose to test the categorization of Te vs DC and Te vs Th because Te and DC belong to the same semantic hyper category of affiliative contact calls and Te and Th are acoustically the most similar (Figure 1). To avoid any confounding effect, the data of one of the females was excluded from the analysis due to an error in the choice of vocalizers: the subject experience the same training day as in Day1 on Day 2 instead of a generalization testing day. Note that the results for this bird on the training and on the non-congruent day (Day 3) was similar to that of the other 7 birds. On Day 1, the subject significantly learn the discrimination

($\log_2(OR_{RC}) = 7.36$ [3.26, 11.46] 95% , Fisher exact test with $p = 1.02 \times 10^{-22}$). On Day 3, the subject significantly classified the calls from the Old vocalizers following the reward contingency rule learned on previous days ($\log_2(OR_{RC}) = 3.80$ [2.44, 5.17] 95% , Fisher exact test with $p = 4.39 \times 10^{-12}$), but was unable to classify the calls from the New vocalizers along the reversed contingency rule. The subject was totally confused and significantly classified these incongruent calls following the previous day rule, resulting in a negative performance ($\log_2(OR_{RC}) = -1.22$ [−2.15, −0.3] 95% , Fisher exact test with $p = 0.010$).

Stimulus Design for Call-Type Discrimination and Categorization.

Since our goals were to assess call-type discrimination and not the identification of particular vocalizers or particular renditions of calls, we used many renditions from many vocalizers for each call-type as stimuli and randomly presented them to the birds as exemplars of call-type. Because of this protocol, birds were forced to use a call categorization strategy to succeed in both tasks.

For Task 1, the stimuli for both the unique Re call-type and the ten other NoRe call-types were chosen randomly prior to each testing day by sampling from the repertoires of twelve vocalizers from our call database. Each acoustic stimulus consisted of a sequence of six or three band-pass filtered (0.25-12 kHz) vocalizations of the same vocalizer and of the same call-type, randomly assigned within a 6s window. More precisely, for the longer Begging sequences and Songs, each stimulus consisted of sequences of three different renditions, while for the other call-types each stimulus consisted of six different renditions. Each stimulus started and ended with a rendition. The 5 or 2 intervals between renditions in a given stimulus were randomly drawn from a uniform distribution. For each call-type and each of the 12 vocalizers, ten such stimuli were generated based on the multiple call renditions from these 12 vocalizers (i.e. ten stimuli per vocalizer). In other words, 12 x 10 = 120 stimuli were generated for each call-type based on the call renditions from 12 distinct vocalizers, totaling 1,320 distinct and new stimuli for the 11 call-types per testing day. Rewarded trials were randomly sampled from 120 stimuli and non-rewarded trials from the remaining 1200 stimuli. Since the average number of trials per day was around 540, on any given day, birds heard very few repeats. On average per day, 43% stimuli were not heard at all, 41% were only heard once, 11% were heard twice and 2.5% were heard three times.

For Task 2, we also used 12 vocalizers for each of the call-type: four of these were used for training on Day 1, another set of four was used to test generalization on the congruent test on Day 2, and another set of four was used to assess categorization on the incongruent test on Day 3. The 6s stimuli were generated as in Task 1.

Statistical Modeling of Behavioral Responses in Operant Conditioning.

In our statistical analyses, we either directly determined if the probability of interruption was different for Re vs NoRe stimuli for a particular bird and rewarded call-type or, we analyzed ensemble results (i.e. across birds, call-types and/or sex) by using generalized mixed effect statistical modeling. Note that to ensure equal representation of each subject and each call-type in the analysis of ensemble results of Task 1, we used the data of the last testing day when the subject took longer than one day (up to three days, see above) to achieve a significant discrimination.

*Comparing two probabilities of interruption*

We used the counts of interrupted stimuli and Fisher exact tests to assess if the bird interruption behavior was statistically different for Re versus NoRe stimuli. For this purpose, we used the python function *fisher_exact()* from *scipy.stats*. The *fisher_exact()* function also provides 95% confidence intervals for the odds ratio of interruption (OR). OR is obtained from the probabilities of interruption of NoRe vs Re stimuli, also using count statistics. We call this OR based on reward contingency (RC) the $OR_{RC}$. The $OR_{RC}$ is a measure of effect size and is defined as:

$$OR_{RC} = \frac{Odds_{int}(NoRe)}{Odds_{int}(Re)}$$

where

$$Odds_{int}(NoRe) = \frac{p_{int}(NoRe)}{1-p_{int}(NoRe)} \text{ and } Odds_{int}(Re) = \frac{p_{int}(Re)}{1-p_{int}(Re)}.$$

with $Odds_{int}(NoRe)$ the odds of interrupting NoRe stimuli and $Odds_{int}(Re)$ the odds of interrupting Re stimuli. Here $p_{int}$, the probability of interruption, is obtained directly from count data. The confidence errors on the $p_{int}$, and thus the corresponding OR confidence errors, are obtained from the binomial distribution. In our results, we expressed this effect size in log2 units, $\log_2(OR_{RC})$. Chance performance yields $OR \approx 1$ or $\log_2 OR \approx 0$. This OR estimate obtained from the Fisher test is used in Fig 2D.

*Analysis of ensemble results in Task 1*

When averaging across birds and call-types, we used the counts of interruption to model the probability of interruption using generalized mixed-effects linear models for binomial distributions. This statistical modeling was performed in R using the *glmer()* function from *lme4*. Using the R compact equation notation, the base statistical model was defined as:

$$cbind(Int, NoInt) \sim Re + ((1 + Re)|Bird)$$

where *Int* and *NoInt* are the counts of interrupted and non-interrupted stimuli, *Re* is a two-level factor describing the reward contingency for each stimulus (*Re*=1 for Re stimuli and *Re*=0 for NoRe stimuli) and *Bird* is the identity of the bird. In this statistical modeling approach, variability across birds is managed by allowing each bird to have its own baseline rate of interruption (random intercept) and its own responsivity to the reward contingency (random slope for *Re*); pseudo-replication is managed by taking into account the number of trials for each bird when estimating probabilities.

To statistically test for the effect of reward contingency, the predictive power of the base model is compared to that of the null model given by:

$$cbind(Int, NoInt) \sim 1 + ((1 + Re)\,|Bird)$$

using a likelihood ratio test (i.e. Chi-square test for differences in deviance).

Similarly to the Fisher exact test, the effect size is measured by the odds ratio (OR) for the reward contingency (RC) in $\log_2$ units, $\log_2(OR_{RC})$.

The coefficients of our statistical model yield estimates of the $OR_{RC}$. More precisely, the base model yields predictions for the probability of interruption $\hat{p}_{int}$ by:

$$\ln \frac{\hat{p}_{int}}{1-\hat{p}_{int}} = \beta_0 + \beta_1 * Re$$

which yields in the case of NoRe stimuli (*Re*=0):

$$\ln \widehat{Odds_{int}(NoRe)} = \ln \frac{\hat{p}_{int}}{1-\hat{p}_{int}} = \beta_0$$

and in the case of Re stimuli (*Re*=1):

$$\ln \widehat{Odds_{int}}(Re) = \ln \frac{\hat{p}_{int}}{1-\hat{p}_{int}} = \beta_0 + \beta_1$$

where $\beta_0$ and $\beta_1$ are the fixed intercept and slope for the *Re* factor . Thus,

$$\log_2(OR_{RC}) = \log_2(\exp(-\beta_1))$$

The estimates of the standard errors (SE) for the model coefficients were used to obtain estimates of the SE for the $\log_2(OR_{RC})$. 95% confidence intervals were obtained from mean ± 2SE.

To test for the effect of sex on discrimination, the base model was compared to the larger model given by:

$$cbind(Int, NoInt) \sim Re * Sex\ +\ ((1\ +\ Re)|Bird)$$

where *Sex* is a two-level factor.

Similarly, to test for the effect of call-type on discrimination, the base model was compared to the larger model given by:

$$cbind(Int, NoInt) \sim Re * Call\ +\ ((1\ +\ Re)|Bird)$$

where *Call* is a 11-level factor describing the nature of the rewarded call-type. Finally, to obtain an effect size ($\log_2(OR_{RC})$) with confidence intervals and p-values for each call-type (shown in Table S1), the base model and null model were also fitted to the subsets of the data partitioned by each call-type (e.g. for the DC call-type, all tests where DC was the rewarded call-type).

*Analysis of ensemble results in Task 2*

For Task 2, as for Task 1, we fitted the data using the same base statistical model:

$$cbind(Int, NoInt) \sim Re\ +\ ((1\ +\ Re)|Bird)$$

We also used the value of the coefficients to determine the $\log_2(OR_{RC})$, and compared it to the null model using a likelihood ratio test to assess whether the $OR_{RC}$ was significantly different from 1. To separately evaluate the performance of discrimination for old and novel stimuli, we fitted these statistical models for the new and old vocalizers separately on each testing day. To statistically determine whether the $OR_{RC}$ was different for old and new vocalizers, we fitted a larger model given by:

$$cbind(Int, NoInt) \sim Re * New\ +\ ((1\ +\ Re)|Bird)$$

where *New* is a two-level factor describing whether the count data is associated to old (New='N') versus new (New='Y') vocalizers. This larger model is compared to the base model using a likelihood ratio test. Finally, we repeated these analyses using only the first trials on the testing days before the birds were able to gain novel information of reward contingency. For a given vocalizer and a given call-type (e.g. Te calls from vocalizer A), these first trials correspond to all trials where renditions of the call-type (Te) of this particular vocalizer (A) were played-back and interrupted by the subject, up to the first trial of the same stimulus type (Te from vocalizer A) that the subject did not interrupt. This last uninterrupted trial is considered as an informative trial because the subject can get knowledge of the rewarding status of this particular vocalizer's call-type: the bird is waiting and can gain knowledge of the outcome (seed access or no seed access) of the particular stimulus (Te from vocalizer A). To investigate which classification rule the subject is applying on the sound stimuli before these first informative trials occur for all combination of vocalizer and call-type, we analyzed the interruption behavior of the subjects on these first "uninformed" trials. To analyze how the birds might update their classification rule with increased knowledge about the rewarding status of stimuli and obtain the learning curves displayed in Figure 3C, we applied the same approach to the following trials in the dataset, grouping them according to the number of informative trials. In other words, the trials used to estimate the interruption behavior after $i$ informative trials correspond, for each call-type and vocalizer, to all trials that happened after the informative trial $i$ until and including the informative trial $i+1$. Counts are aggregated across vocalizers to estimate the probabilities. Depending on individual behavioral differences, the number of informative trials a bird might accumulate over the course of a testing day is variable between subjects. As a result, not all subjects went up to 11 informative trials for at least one combination of vocalizer and call-type.

<u>Quantifying correct classification and systematic misclassification of each Call-type in Task 1</u>

To quantify and represent how each call-type is correctly classified as a rewarded or non-rewarded stimulus by the bird, we calculated the odds ratio of correct behavioral classification $OR_{Beh}$ for each call-type (Figure 2B, and 2C). The rationale is that the subject classifies a stimulus as non-rewarded by interrupting it and classifies a stimulus as rewarded by refraining to interrupt it. We can then calculate for each call-type the odds of correct behavioral response which is the odds of interrupting if the call-type is a non-rewarded stimulus and the odds of *not* interrupting if the call-type in a rewarded stimulus. These odds are then compared in the odds ratio $OR_{Beh}$to the odds of incorrect behavioral response for the opposite stimulus class.

In the case where the call-type $i$ is rewarded, the correct behavioral response is refraining to interrupt $i$ stimuli. So the degree with which this call-type is correctly classified as rewarded is measured by comparing the odds of *not* interrupting $i$ stimuli (correct response for these stimuli) to the baseline (average) odds of *not* interrupting all the other call-types $j \neq i$ (incorrect response for these non-rewarded stimuli).

The odds of *not* interrupting the call-type $i$ given it is the rewarded call-type is given by:

$$Odds_{noint}(i|i) = \frac{1-p_{int}(i|i)}{p_{int}(i|i)} \quad [1]$$

The odds of *not* interrupting any other call-type $j \neq i$ given $i$ is the rewarded call-type is given by:

$$Odds_{noint}(j \neq i|i) = \frac{1-(\sum_{j\neq i} p_{int}(j|i)/(n-1))}{(\sum_{j\neq i} p_{int}(j|i)/(n-1))} \quad [2]$$

$OR_{Beh}(i|i)$ is then given by:

$$OR_{Beh}(i|i) = \frac{Odds_{noint}(i|i)}{Odds_{noint}(j\neq i|i)} \quad [3]$$

Note that $OR_{Beh}(i|i)$ provides a separate estimate of the $OR_{RC}$ for each call-type that takes into account the varying presentation rate of non-rewarded call-types.

For each of the other non-rewarded call-types $j$, the correct behavioral response is to interrupt $j$ stimuli. So the degree with which this call-type $j$ is correctly classified as non-rewarded is measured by comparing the odds of interrupting $j$ stimuli (correct response for these stimuli) to the baseline odds of interrupting the rewarded call-type $i \neq j$ (incorrect response for these rewarded stimuli).

The odds of interrupting the call-type $j$ given $i$ is the rewarded call-type is given by:

$$Odds_{int}(j|i) = \frac{p_{int}(j|i)}{1-p_{int}(j|i)} \quad [4]$$

The odds of interrupting the call-type $i$ given it is the rewarded call-type is given by:

$$Odds_{int}(i|i) = \frac{p_{int}(i|i)}{1-p_{int}(i|i)} \quad [5]$$

$OR_{Beh}(j|i)$ is then given by:

$$OR_{Beh}(j|i) = \frac{Odds_{int}(j|i)}{Odds_{int}(i|i)} \quad [6]$$

To obtain the learning curves in Figure 2B, 2C, S1 and S2, we calculated $OR_{Beh}$ for data binned in sets of consecutive trials that included ten rewarded trials. At each time bin, we tested if the odds ratio was different from 1 using a Fisher exact test.

For each subject, these values of correct behavioral classification $OR_{Beh}$ are gathered in a matrix and quantify the discrimination between each pair of call types (right panels of Figure 2B and 2C). The diagonal terms $OR_{Beh}(i|i)$ provide a separate estimate of the $OR_{RC}$ for each call type. The off-diagonal terms $OR_{Beh}(j|i)$ quantify the correct classification (blue color) of each call-type (columns) as a non-rewarded stimulus depending on what was the rewarded call-type on that particular test day (rows). $OR_{Beh}$ equals to 1 when the subject is performing at chance level and the degree to which $OR_{Beh}$ is greater than 1 quantifies the discrimination performance. If $OR_{Beh}$ is significantly less than 1, then the bird is systematically misclassifying the call-type. In the matrix of Figure 2B and 2C, we use a graded red color to show call-types classified as rewarded and a graded blue color to show call-types classified as non-rewarded. The off-diagonal terms reveal systematic errors (misclassification) when a non-rewarded call-type is systematically classified as the rewarded call-type (red color): the probability of interruption for that non-rewarded call-type is less than the one observed for the rewarded call-type.

Visualizing correct classification and systematic misclassification in Task 2

For task 2, our goal was to assess how birds are generalizing the reward contingency rule to new vocalizers on Day 2 and how they are affected by the inversion of reward contingency on Day 3. We expected that birds could be confused on Day 3, since we flipped the reward contingency for New vocalizers. To clearly visualize which calls are classified as Rewarded vs Non-Rewarded by the birds, we estimated the odds ratio of classifying calls as rewarded by grouping calls according to each combination of Etho call-type (Te, Th or DC) and vocalizer type (Old or New). There were a total of six call classes: Te-New, Te-Old, DC-New, DC-Old, Th-New, Th-Old. For each class, we calculated the odds that the bird would classify this class of calls as rewarded $Odds_{Re}(Class)$ and compared this odds to the average odds of classifying any calls (both Re and NoRe) from the Old vocalizers as rewarded $\overline{Odds_{Re}(Old)}$. This latter average odds was estimated across the entire day and defined a steady state baseline for the classification of calls that the subject knew very well. The ratio of these odds, called Odds Ratio for Classification, $OR_{Class}$, is greater than 1 when the stimulus class is interrupted less than average, corresponding to classifying the stimulus class as rewarded, and is less than 1 when the stimulus class is interrupted more than average, corresponding to classifying the stimulus class as non-rewarded. To obtain time-varying $OR_{Class}(t)$, the $Odds_{Re}(Class)$ was estimated for sliding time windows composed of 10 trials from each stimulus class $Odds_{Re}(Class, t)$. Similar to equation [1], the odds of classifying the stimulus class as a rewarded stimulus at time $t$ is equal to the odds of *not* interrupting that stimulus class:

$$Odds_{Re}(Class, t) = Odds_{noint}(Class, t) = \frac{1 - p_{int}(Class, t)}{p_{int}(Class, t)}$$

with $p_{int}(Class, t)$ the probability of interrupting all trials playing stimulus class $Class$ in the time window $t$.

Similarly, the odds baseline of classifying any calls from the Old vocalizers as rewarded stimuli is equal to the odds of *not* interrupting the calls from the Old vocalizers:

$$\overline{Odds_{Re}(Old)} = \overline{Odds_{noint}(Old)} = \frac{1 - \overline{p_{int}(Old)}}{\overline{p_{int}(Old)}}$$

with $\overline{p_{int}(Old)}$ the time average probability of interrupting all trials playing calls from *Old* vocalizers.

$OR_{Class}(t)$ can then be written in equation form as:

$$OR_{Class}(t) = \frac{Odds_{Re}(Class, t)}{\overline{Odds_{Re}(Old)}} = \frac{\dfrac{1 - p_{int}(Class, t)}{p_{int}(Class, t)}}{\dfrac{1 - \overline{p_{int}(Old)}}{\overline{p_{int}(Old)}}}$$

where the bar on top of $\overline{p_{int}(Old)}$ is used to indicate the average across time.

$OR_{Class}(t)$ is shown in Figs 3A and 3B for two example subjects and for illustrative purposes. Values are plotted on a logarithmic scale as the absolute value of $\log_2 OR_{Class}(t)$ and the colored axis gives the sign (red for positive, blue for negative).

Etho call-types acoustical analyses and acoustic confusion matrices

A detailed analysis of the acoustic characteristics each Etho call-type was performed in (*7*). In the present study, we selected the optimal acoustic representations of sounds for the classification of calls into Etho call-types, to investigate how behavioral discrimination could be explained by the acoustic characteristics of calls. In (*7*), we used three types of supervised classifiers operating on four distinct acoustic feature spaces (input to the classifier) to assess the degree with which the calls could be discriminated into the Etho call-types. The performances of classifiers were quantified by cross-validation on calls from birds (entire repretoires from vocalizers) that were left-out of the training set. These cross-validated performances were very similar between the three types of supervised classifiers and here, we chose to use the Linear Discriminant Analysis (LDA). Among the four feature spaces investigated in (*7*) to represent the sounds, the spectrograms and Predefined Acoustic Features (PAF) gave the highest levels of Etho call-type discrimation. For the results presented here, we used either spectrographic values or the PAF as a feature vector to represent each sound depending on whether a linear representation was desirable or not for the analysis. The PAF included 20 features: five features describing the fundamental (saliency, average F0, max F0, min F0, coefficient of variation of F0), eight features describing the spectral envelope (mean, std, skew, kurtosis, entropy, Q1, Q2, Q3), five features describing the temporal envelope (mean, std, skew, kurtosis, entropy) and two features describing the intensity (rms, max amplitude). The spectrographic features were obtained by first centering each call in a 350 ms window using the mean of the temporal envelope. For calls shorter than 350 ms, the edges of the window were padded with zeros. The spectrogram of these cut sounds was estimated using gaussian windows of frequency bandwidth of 52 Hz which yielded a matrix of 231 frequency bands and 357 points in time. To reduce the dimensionality of this feature space, we applied a principal component analysis to all spectrograms. Using cross-validation in our discriminant analyses, we found that the maximum performance was obtained with 40 principal components (PCs). Thus, we used these 40 spectrogram PCs as a spectrographic feature vector to represent each sound.

The 20 PAFs and 40 spectrographic features were used to create 2-dimensional visualizations of the acoustic space spanned by the different Etho call-types (Fig 1A and S1). These visualizations were obtained by applying the uniform manifold approximation and projections (UMAP) implemented in the Python library *umap.* The UMAP is a non-linear embedding of a higher dimensional space into a lower dimensional manifold based on local distances between data points.

To estimate the degree of acoustic similarity of calls belonging to the same Etho call-type (Fig. S1), we first measured the pair-wise distances between calls in each of the two UMAPs (spectrogram derived and PAF derived). We then clustered the calls by independently applying unsupervised hierarchical clustering to each set of distances (*linkage* function from the python library *scipy.cluster*). The adjusted Rand index was calculated to quantify the match between each of the unsupervised clustering obtained that way, with the grouping of calls into Etho call-types.

To compare the algorithmic classification of calls to the classification performed by the birds in the operant task 1, we chose to operate in a linear acoustic space to measure the performance of acoustic classification. The spectrographic features in combination with the LDA yield a euclidian acoustic space where distances between calls can be interpreted as distances in acoustic (physical) space. The posterior probabilities of the LDA applied to spectrographic features were used to generate acoustic confusion matrices showing the correct classification (diagonal) and the systematic miss-classifications (off-diagonal) of calls along Etho call-types. One acoustic confusion matrix was generated for each bird in the call database (n=45) using the leave-one-bird-out cross-validation approach. To account for the different number of examplars of each Etho call-type between the different birds of the call database, these acoustic confusion matrices recorded the number of calls classified in each cell of the confusion matrix (i.e. a pair of actual and predicted call-types) based on the maximum posterior probability (winner take all procedure). These 45 count-based acoustic confusion matrices were used in a bootstrap procedure to estimate the variability in the calculation of acoustic distances between Etho call-type and in the quantification of the semantic magnet effect described below (see *Acoustics and Semantic Hyperclustering*). In brief, each bootstrap loop generates a bootstrapped posterior conditional probability matrix by summing 45 random samples of these count-based matrices and dividing each cell by the row marginals. The orginal analyses in (*7*) were performed using custom MATLAB code. That code has now been updated and exported to Python and is publicly available at https://github.com/theunissenlab/soundsig with tutorials at https://github.com/theunissenlab/BioSoundTutorial

Generating an Odds Ratio Matrix from the Acoustic Confusion Matrix

In our acoustical analyses, a supervised classifier (LDA) trained on the spectrogram and tested on cross-validated data returns the bootstrapped posterior conditional probability of a call $c$ belonging to call-type $i$ to be identified at $j$: $p_{class}(c = j|i)$. To translate our acoustic confusion matrices obtained from the classifier into acoustic OR matrices, $OR_{Ac}$, that could be compared to $OR_{Beh}$ matrices obtained from birds' behavior, we used the same logic and equations [1-6] shown above with a few twists.

For the diagonal terms of the $OR_{Ac}$ matrix, $p_{class}(c = i|i)$ is the probability of correct classification, while the probability of incorrect classification is the probability of classifying a call $c$ belonging to call-type $i$ as any $j$, $p_{class}(c = j|i)_{k\in\{j\},i\notin\{j\}}$ . If you look back at the quantification of behavioral discrimination (previous methods section), $p_{class}(c = i|i)$ is equivalent to $1 - p_{int}(i|i)$ and both are the probability of correct classification for the LDA and correct decision for behavior. Similarly to equation 1 above we can then define the equivalent of $Odds_{noint}(i|i)$ for the LDA as:

$$Odds(c = i|i) = \frac{p_{class}(c=i|i)}{1-p_{class}(c=i|i)} \quad [1b]$$

Then similarly to equation [2] we can use $p_{class}(c = k|i)_{k\in\{j\},i\notin\{j\}}$ to calculate the odds of incorrect classification:

$$Odds(c = k|i)_{k\in\{j\},i\notin\{j\}} = \frac{p_{class}(c=k|i)_{k\in\{j\},i\notin\{j\}}}{1-p_{class}(c=k|i)_{k\in\{j\},i\notin\{j\}}} = \frac{(\sum_{j\neq i} p_{class}(c=j|i)/(n-1))}{1-(\sum_{j\neq i} p_{class}(c=j|i)/(n-1))} \quad [2b]$$

The ratio of these odds then yields the odds ratio of correct algorithmic classification of call $c$ as call-type $i$ based on acoustic (similar to equation [3]):

$$OR_{Ac}(i,i) = \frac{Odds(c=i|i)}{Odds(c=k|i)_{k\in\{j\},i\notin\{j\}}} \quad [3b]$$

For the off-diagonal terms, we compare the odds of avoiding a misclassification (the odds of avoiding classifying call $c$ as call-type $j$ given it belongs to call-type $i$, $Odds(c \neq j|i)$) to the odds of general misclassification (odds of classifying call $c$ as *not* call-type $i$ given it belongs to call-type $i$, $Odds(c \neq i|i)$). $Odds(c \neq j|i)$ is similar to the odds quantifying the correct behavioral classification in the case where the stimulus does not belong to the rewarded call-type (see equation [4] above). The probability of correct behavioral response $p_{int}(j|i)$ is similar to the probability of correct algorithmic classification which is the probability of not classifying call $c$ as call-type $j$ given it belongs to call-type $i$: $1 - p_{class}(c = j|i)$.

Thus, similarly to equation [4], we can write the odds of correct algorithmic decision as:

$$Odds(c \neq j|i) = \frac{1-p_{class}(c=j|i)}{p_{class}(c=j|i)} \quad [4b]$$

The odds of general misclassification, $Odds(c \neq i|i)$), is equivalent to the odds quantifying the behavioral misclassification of the rewarded call-type (see equation [5] above). The probability of incorrect behavioral response (interrupting a rewarded call-type) $p_{int}(i|i)$ is similar to the probability of incorrect algorithmic classification which is the probability of not classifying call $c$ as call-type $i$ given it belongs to call-type $i$: $1 - p_{class}(c = i|i)$.

Thus, similarly to equation [5], we can write the odds of incorrect algorithmic decision as:

$$Odds(c \neq i|i) = \frac{1-p_{class}(c=i|i)}{p_{class}(c=i|i)} \quad [5b]$$

Similar to equation [6], $OR_{AC}(j|i)$ is then given by:

$$OR_{Ac}(i,j) = \frac{Odds(c\neq j|i)}{Odds(c\neq i|i)} \quad [6b]$$

A large (significant) positive diagonal term quantifies the classifier discrimination for that particular row call-type. A large (significant) off-diagonal term shows that the row call-type has a low probability of being misclassified as the column call-type. The OR matrix, $OR_{Ac}$, and the acoustic confusion matrix provide the same information.

The off-diagonal terms of the $OR_{Beh}$ and $OR_{Ac}$ matrices in log units are then used as measures of "distance" between Etho call-types and quantify behavioral and acoustic separation based on the observed systematic misclassifications.

Generating Spatial Maps from the Odds ratio Matrices

The off-diagonal terms in the $OR_{Beh}$ and $OR_{Ac}$ matrices were used as distances in a Multi Dimensional Scaling analysis to obtain a map of the Etho call-types in respectively the perceptual space and the acoustic space (Figure 4B). Following a bootstrap procedure described below, the matrices obtained from several bird subjects in Task 1 or several bird vocalizers in the call database were first averaged to generate one matrix for each space. Then these two matrices followed the same procedure: the i,j and j,i off-diagonal terms were averaged together to generate a triangular matrix and the minimum diagonal term was subtracted from all off-diagonal terms to generate positive distances, with the assumption that a distance of a call to itself should be zero. Multi-dimensional scaling was done with the MDS() function in the Python library *sklearn.* Data was projected to 2 dimensions. Similar results were obtained in 3D (data not shown). To line up the 2D MDS spaces obtained from the acoustic and behavioral data we applied a Procrustes scaling and rotation using the procrustes() function from the Python library *scipy.spatial*.

Quantifying the clustering of Etho call-types into hyper-categories

For each of the MDS acoustical and behavioral spaces, the Q factor quantifies the clustering of Etho call-types into hyper-categories by measuring how close each Etho call-type is on average to the semantic hyper-category it belongs to, as compared to other semantic hyper-categories. It is calculated as the ratio of the between to within square distances from the position of each call-type to the centroid of the semantic hyper-categories:

$$Q = \frac{\sum_{b \neq w}^{n_h - 1} \sum_{j}^{n_c} \left\| \overrightarrow{x_j} - \overrightarrow{y_b} \right\|^2 / (n_c * (n_h - 1))}{\sum_{j}^{n_c} \left\| \overrightarrow{x_j} - \overrightarrow{y_w} \right\|^2 / n_c}$$

where $n_c = 11$ is the number of call-types; $n_h$ is the number of semantic hyper-categories, $\overrightarrow{x_j}$ is the position in the 2d MDS space of call-type $j$; $\overrightarrow{y_b}$ is the position of the centroid of all semantic hyper-categories that don't include the call-type, $j$ (there are $n_h - 1$); and $\overrightarrow{y_w}$ is the position of the centroid of the unique semantic hyper-category that includes the call type $i$. Just like an F statistic, the size of the Q value beyond 1 assesses the degree of clustering of the semantic hyper-categories. Separate Q values were obtained for the acoustic space, $Q_{Ac}$, and behavioral space, $Q_{Beh}$ .

Resampling statistical analyses for spatial maps and the Q factor

To assess the robustness of our generated spatial maps and the semantic hyper-category clustering we used resampling techniques. We used a single bootstrap to randomly sample the 45 birds used in the acoustic analyses (as providers of call exemplars) and the 12 birds used in the behavioral analysis (as subjects discriminating call-types). For each iteration of the bootstrap (1000 total), we randomly chose (with replacement) 45 count-based acoustic confusion matrices $OR_{AC}$ and 12 count based behavioral confusion matrices $OR_{Beh}$. From these random samples, average acoustic and confusion matrices were generated and their off-diagonal terms used to generate the corresponding acoustic and spatial maps (described above and shown in Figure 4B). Q values were also estimated for each bootstrap iteration. In addition, within each bootstrap, we performed 1,000 permutations of hyper-category definitions by randomly choosing partitions of call-types into one group of 3, three groups of 2, and two groups of 1 call-type (corresponding to the group numeracy of the actual hyper-categories). 1,000 permuted Q values were obtained for each of the bootstrap samples. With this nested resampling procedure, we were able to obtain 1,000 actual Q values by bootstrap and 100,000 permuted Q values for both the acoustic and behavioral spaces (shown in Figure 4D). To assess whether the actual Q value had a value greater than expected by chance (as obtained in random hyper-categories), we calculated the number of times the actual Q value obtained in each bootstrap iteration was larger than the 1,000 permutations. Finally, to assess whether the behavioral Q value was greater than the acoustical Q value, we counted the number of bootstrap iterations for which we found $Q_{Beh} > Q_{Ac}$. This relationship was found 996 times out of 1000, yielding a p value of $p = 0.004$.

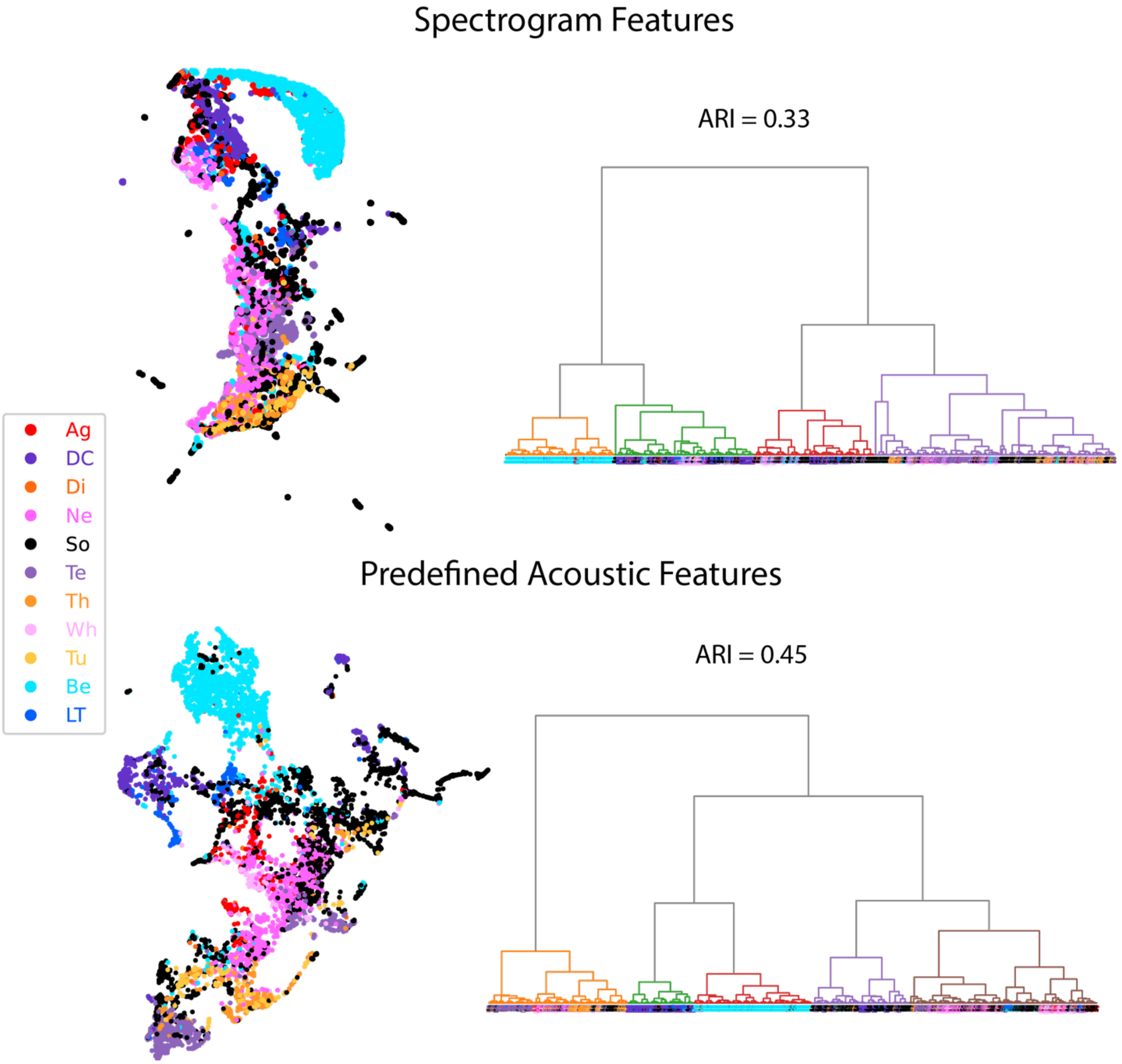


**Fig. S1.** The scatter plots on the left panels show the UMAP 2d-projections of all calls in the database using spectrographic features (top) or predefined acoustic features (bottom, same as in Fig. 1A). Each call is labelled according to the Etho call-type it belongs to (dot color). The pair-wise distances in these projections were used in unsupervised hierarchical clustering to obtain the dendrograms shown on the right panels. The leaves of these dendrograms are also colored according to the Etho call-type to visualize the match between the unsupervised clustering based on acoustics and the grouping based on ethological assessments. The adjusted Rand index (ARI) quantifies this match.

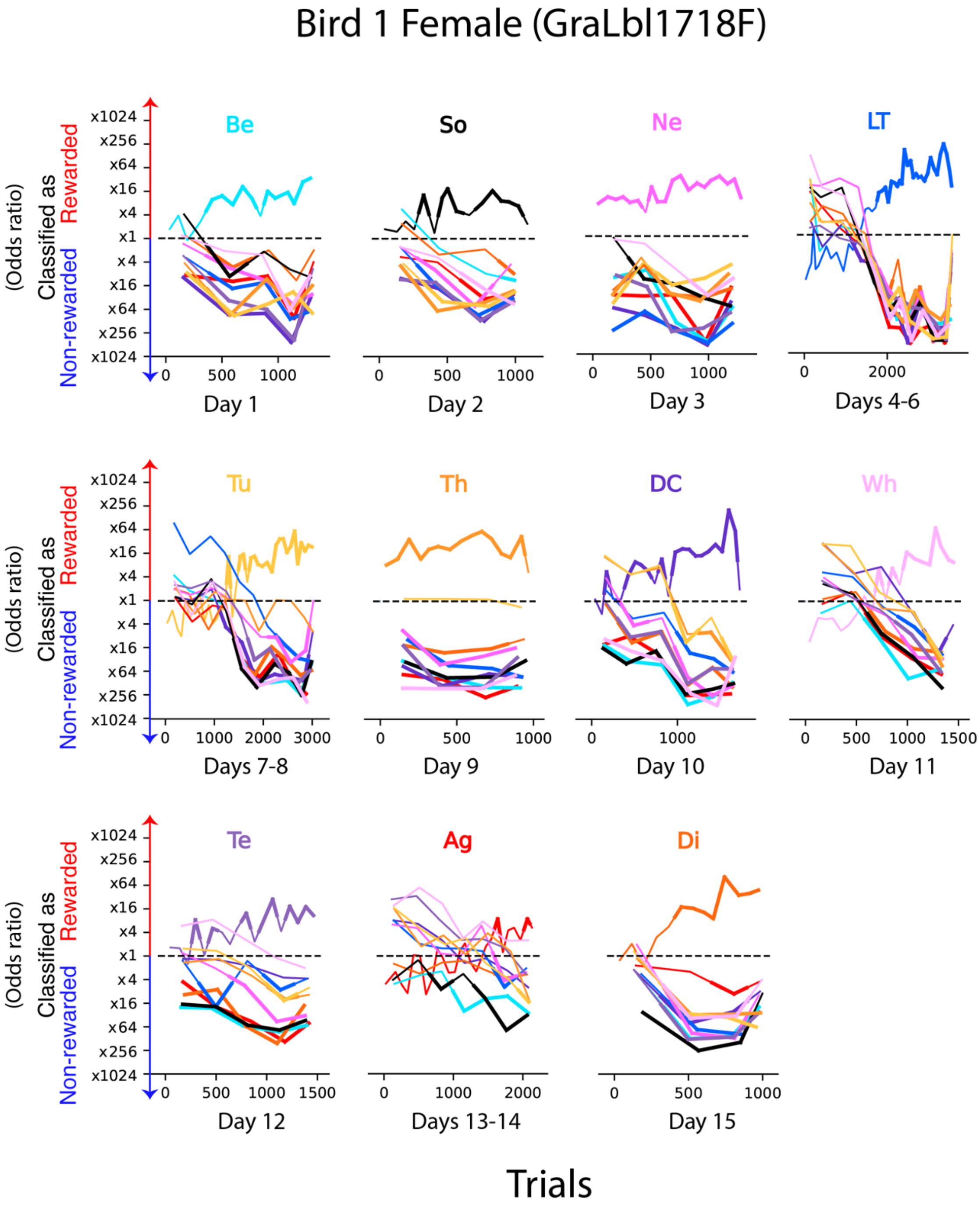


**Fig. S2.**

Performances of the female bird shown in Figure 2B in the Etho call-type discrimination task. For a given call-type test, birds have up to three consecutive days to identify the rewarded call-type and show their discrimination performance. All tests are shown in chronological order. The learning curves show for each Etho call-type (line color) the average bird performance at classifying calls as rewarded or non-rewarded stimuli. The performance is quantified by the odds ratio of correct behavioral classification $OR_{Beh}$ (see legend of Figure 2B and Methods).

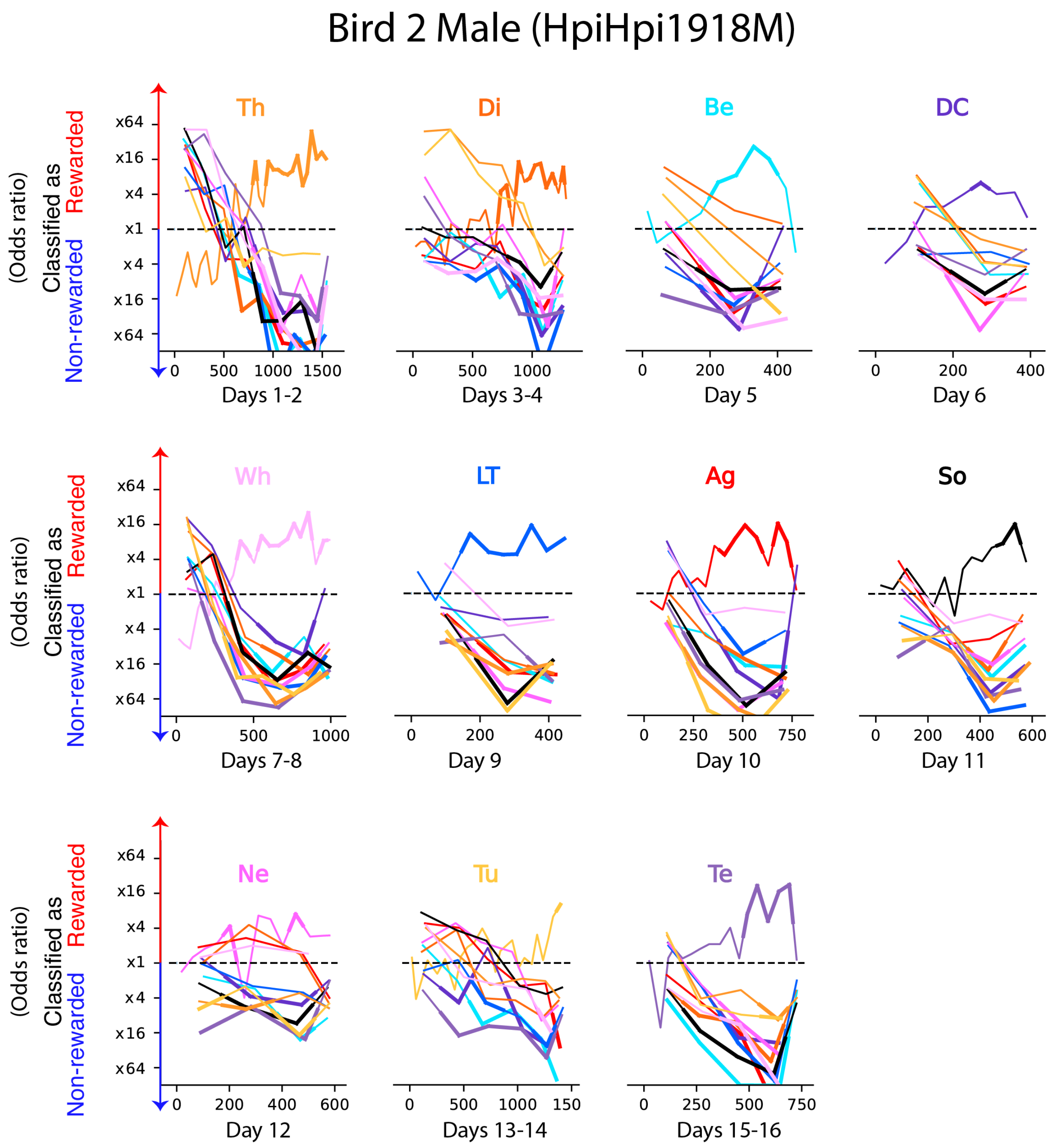


**Fig. S3.**

Same as Figure S2 for the male bird shown in Figure 2C.

| Call-type | Effect Size ($Log_2(OR)$) | 2 SE of Effect Size |
|---|---|---|
| So | 3.42 | 0.827 |
| DC | 3.33 | 0.009 |
| LT | 3.00 | 0.982 |
| Te | 3.53 | 0.011 |
| Tu | 2.87 | 0.948 |
| Th | 3.37 | 0.011 |
| Ag | 2.16 | 1.014 |
| Di | 2.76 | 0.801 |
| Wh | 3.03 | 1.126 |
| Ne | 2.31 | 0.819 |
| Be | 2.77 | 0.771 |

**Table S1.**

Zebra finches' discrimination performance for one call-type vs all others (Task 1) also plotted in Figure 2D. The effect size is quantified by the $\log_2$ of the Odds Ratio for the reward contingency ($OR_{RC}$). Chance level is $\log_2(OR_{RC}) = 0$. All call-types are significantly discriminated from all others (mixed-effect logistic regression; likelihood ratio test, all tests $p < 0.001$).

| Learning rule on Day 1 | Rewarded call-type | Non rewarded call-type | Number of birds |
|---|---|---|---|
| Te vs DC | DC | Te | 2 |
| | Te | DC | 2 |
| Te vs Th | Th | Te | 2 |
| | Te | Th | 1 |

**Table S2.**

Number of birds tested in each condition in Task 2.
